# Multi-dimensional structure function relationships in human β-cardiac myosin from population scale genetic variation

**DOI:** 10.1101/039321

**Authors:** Julian R. Homburger, Eric M. Green, Colleen Caleshu, Margaret Sunitha, Rebecca Taylor, Kathleen M. Ruppel, Raghu Metpally, SHaRe Investigators, Steven D. Colan, Michelle Michels, Sharlene Day, Iacopo Olivotto, Carlos D. Bustamante, Frederick Dewey, Carolyn Y. Ho, James A. Spudich, Euan A. Ashley

## Abstract

Myosin motors are the fundamental force-generating element of muscle contraction. Variation in the human β-cardiac myosin gene (*MYH7*) can lead to hypertrophic cardiomyopathy (HCM), a heritable disease characterized by cardiac hypertrophy, heart failure, and sudden cardiac death. How specific myosin variants alter motor function or clinical expression of disease remains incompletely understood. Here, we combine structural models of myosin from multiple stages of its chemomechanical cycle, exome sequencing data from population cohorts of 60,706 and 42,930 individuals, and genetic and phenotypic data from 2,913 HCM patients to elucidate novel structure-function relationships within β-cardiac myosin. We first developed computational models of the human β-cardiac myosin protein before and after the myosin power stroke. Then, using a spatial scan statistic modified to analyze genetic variation in protein three-dimensional space, we found significant enrichment of disease-associated variants in the converter, a kinetic domain that transduces force from the catalytic domain to the lever arm to accomplish the power stroke. Focusing our analysis on surface-exposed residues, we identified another region enriched for disease-associated variants that contains both the converter domain and residues on a single flat surface on the myosin head described as a myosin mesa. This surface is prominent in the pre-stroke model, but substantially reduced in size following the power stroke. Notably, HCM patients with variants in the enriched regions have earlier presentation and worse outcome than those with variants in other regions. In summary, this study provides a model for the combination of protein structure, large-scale genetic sequencing and detailed phenotypic data to reveal insight into time-shifted protein structures and genetic disease.

---

Myosin motors are molecular machines responsible for converting chemical energy into the mechanical force necessary for cell division, directed cell migration, vesicle trafficking and muscle contraction^1^. Efforts to understand structure/function relationships within myosin have been ongoing for more than fifty years, and incorporate structural biology, *in vitro* biophysical and biochemical analyses, and studies in model systems from Dictyostelium to mouse^2,3^. Variants in myosin genes cause several skeletal and cardiac myopathies^4^ including hypertrophic cardiomyopathy (HCM), a genetic disease of the heart muscle characterized by an asymmetric thickening of the ventricular walls and a decrease in the ventricular chamber size. Clinically, HCM can be associated with arrhythmia, heart failure or sudden death^5^. Except in cases with large kindreds, relationships between genotype and disease expression have been challenging to establish due to the absence of large scale genetic population data and lack of multi-center sharing of patient genetic and clinical data^6,7^. Further, limited understanding of three dimensional protein structural dynamics has prevented the extension of inference from genetic variation beyond the single linear dimension of genomic DNA sequence.

Recently, advances in next-generation sequencing technology have enabled the assembly of large datasets of human genetic variation in both unselected and disease-affected populations. Comparative analysis of these cohorts enables within-gene inference of constraint - a measure of population tolerance to variation that can reveal insight into critical functional residues. The *MYH7* gene, encoding the β-cardiac myosin implicated in hypertrophic cardiomyopathy, is highly constrained for missense and loss of function variants^8,9^. Studies of regional tolerance within *MYH7* report conflicting results and suffer from small samples sizes or a lack of a control cohort^6,10,11^. We hypothesized that assessing regional genetic tolerance in the context of time-shifted three-dimensional structures would reveal novel insights into structure-function relationships in *MYH7*.

The Sarcomeric Human Cardiomyopathy Registry (SHaRe) was established as an international consortium of HCM investigators and currently contains detailed longitudinal clinical data on 2,913 HCM patients who have undergone clinical genetic testing^12^. The Exome Aggregation Consortium (ExAC) is a publicly available database of exome sequences from 60,706 unselected individuals^13^. We compared the prevalence of missense variants in *MYH7* within these cohorts. We found 193 unique missense variants (in 476 patients with *MYH7* variants) in the HCM cohort and 454 unique missense variants in the ExAC database. In both cases, observed missense variants were very rare (Supplementary Figure S1), consistent with previous reports of constraint within the *MYH7* gene^8,14^. Although both disease and population variants are non-uniformly distributed throughout the gene, there is a significant difference in the linear distribution of rare variants between these cohorts (KS p = 5.0x10^−11^, Supplementary Figure S1). Disease-associated missense variants are concentrated in the catalytic globular domain and the coiled coil S2, consistent with previous results^6^. Even within these domains, however, distributions of disease and population variants are not the same (KS p = 0.003). These results suggest that the likelihood of *MYH7* variants causing disease is in part due to their location within the gene.

Since molecular motors act in three-dimensional space, we sought a method to investigate patterns of genetic tolerance in the folded structure of human β-cardiac myosin protein. We used multi-template homology modeling of other myosin proteins in the pre- and post-stroke states to build three-dimensional models of human β-cardiac myosin containing the human ventricular light chains (Fig. 1 and Methods). These models represent two distinct phases of the actin-activated myosin chemomechanical cycle. Four fundamental regions of the myosin motor domain are included: the actin-binding site (Fig. 1, green residues), the ATP-binding pocket (red), the converter domain (blue), and the light chain binding region or lever arm. In the pre-stroke state, the converter aligns with a relatively flat surface of the myosin head described as the myosin mesa. Based on its size (>20 nm^2^), flat topology and high degree of evolutionary conservation, this feature has been proposed as an interaction site for intra- or intermolecular binding^15^. Following the force-producing lever arm stroke of a myosin head, the motor is in its post-stroke state (Fig. 1a) and the mesa falls out of alignment with the converter domain.

**Fig 1.**
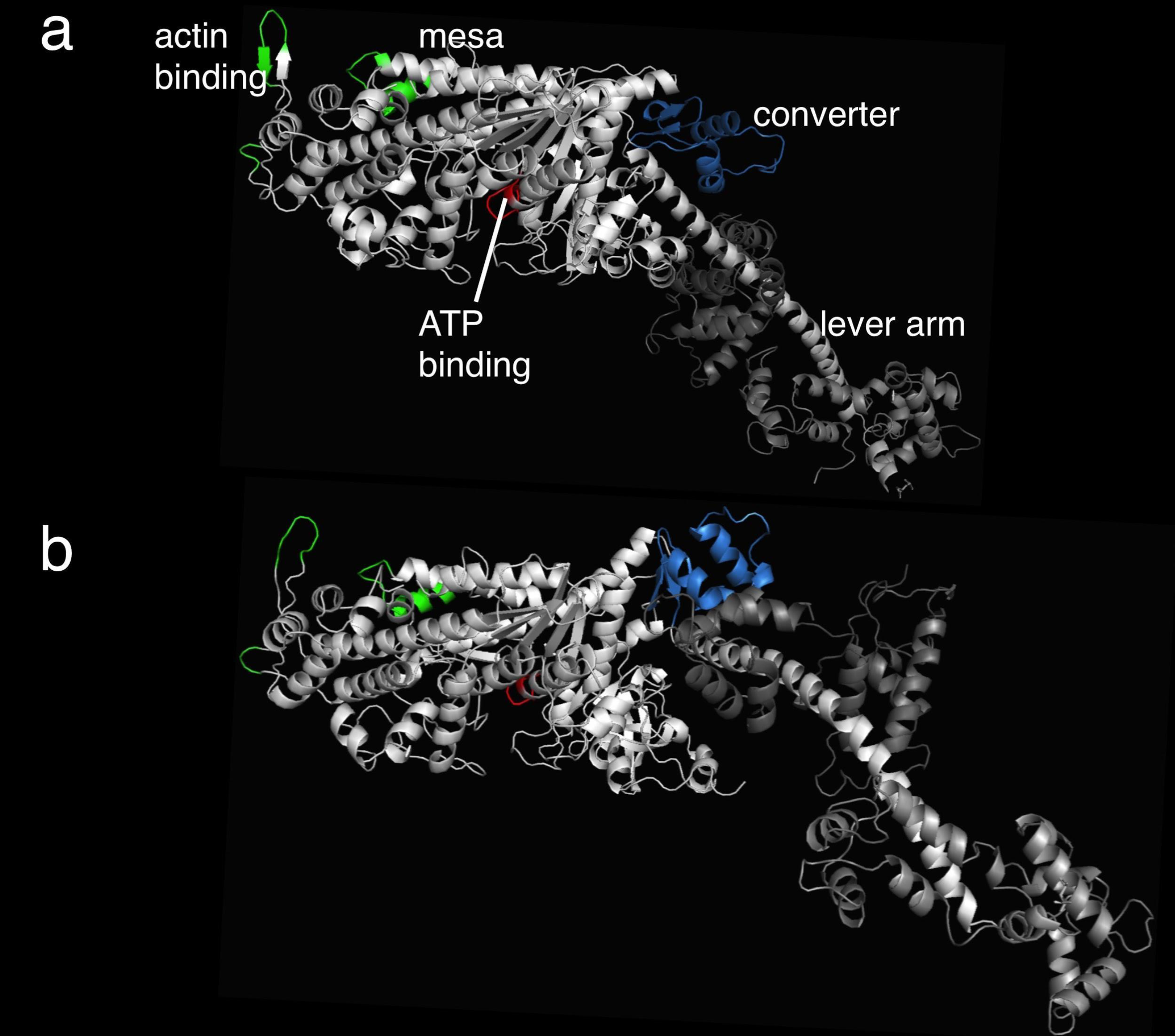
Structural models of the human β-cardiac post- and pre-stroke obtained by integrating data from solved crystal structures of homologous models. (a) A side view of myosin S1, with the relatively flat mesa at the top, in the post-stroke state with important functional domains labeled: the actin-binding site (green residues), the ATP-binding pocket (red), the converter (blue), and the light chain binding region or lever arm. The converter and its associated lever arm is behind the plane of the figure and below the level of the mesa. (b) The myosin S1 in the pre-stroke state. Small changes within the globular head region with ADP and Pi in the nucleotide pocket result in a large ~70° rotation of the converter and lever arm. The converter is moved forward and up compared to the post-stroke structure, and the lever arm is projecting forward out of the plane of the image. The distance traversed by the C-terminal end of the lever arm is ~ 10 nm, the stroke size of the motor.

To prioritize three-dimensional structural regions of interest, we applied a modified version of the spatial scan statistic^16,17^ to the pre-stroke and post-stroke models of β-cardiac myosin S1. This statistic searches for spherical regions with an increased proportion of genetic variants in disease compared to control cohorts. In the myosin pre-stroke model, we find a striking increase in the proportion of disease-associated missense variation in a 15 Å-sphere centered on residue 736 (p = 0.001) (Fig. 2a). This region, covering much of the converter domain, contains 17 missense variants observed in disease and no missense variants observed in control data (Fig. 2c). Using the post-stroke model of β-cardiac myosin, we again observed enrichment of disease-associated variants in the converter domain (p < 0.001) centered on residue 733 (Supplementary Figure S2). Enrichment of disease-associated variants in both the pre- and post-stroke states persists when including only variants formally classified as pathogenic or likely pathogenic, and when limited only to individuals of European descent (Supplementary Text 3). During systolic contraction of the heart, the converter domain serves the critical function of transducing force by swinging about 70° from its pre-stroke position (Fig. 1b, lever arm projecting outward). Variants in the converter domain have been shown to alter muscle power output and kinetics^18,19^ and have been associated with worse outcomes in HCM^20–22^. Our data provide a complementary line of evidence that variants in the β-cardiac myosin converter domain are poorly tolerated and prone to development of HCM.

**Fig 2.**
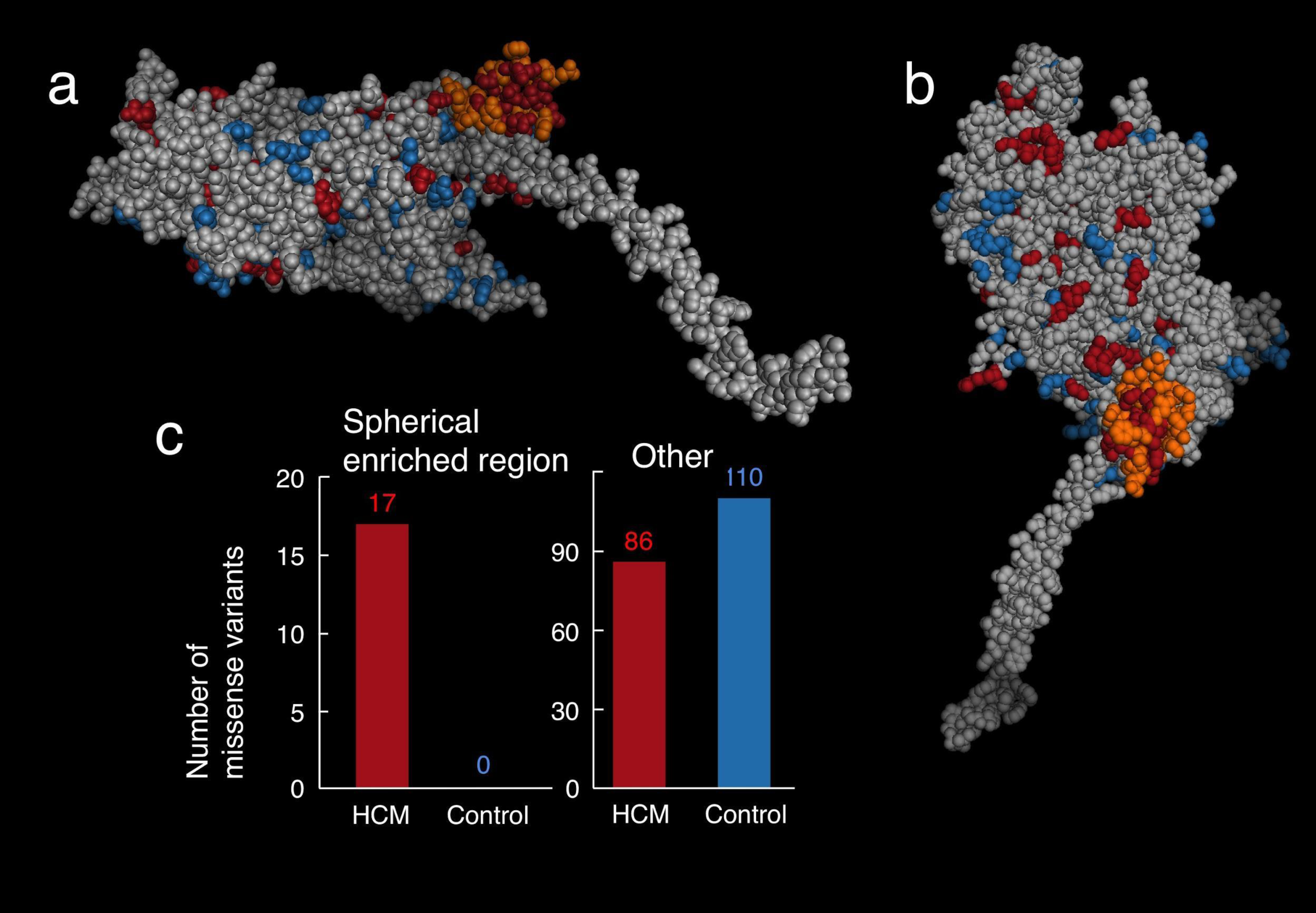
The spatial scan statistic identifies a spherical region of the converter domain with an increased proportion of HCM-associated variants. (a) The same side view of the pre-stroke S1 motor domain as shown Figure 1b, without the light chains attached to the α-helical light chain binding region of the S1. The orange residues define a sphere of residues in the motor domain that is the only region significantly-enriched for HCM variants. The S1 residues are colored as follows: orange - region enriched for HCM variants, blue - missense variants seen only in Exac, red - missense HCM variants seen in SHaRe, light grey - all other residues. (b). The enriched region in the pre-stroke model of myosin, from a different perspective. The view is directly down onto the surface. Coloring is as in (a). (c). The number of HCM-variants (SHaRe) and non-disease associated (ExAC) variants identified in the spherical enriched region (left) and in the sum of all other parts of the myosin (right).

To replicate these results, we sought an independent source of disease-associated and population genetic variation. We curated publications from medical centers not yet affiliated with the SHaRe registry (Supplementary Text 1). We compared this set of 231 missense variants with 430 missense variants found in 42,930 exomes from unselected individuals in the Geisinger Health System sequenced by the Regeneron Genetics Center (DiscovEHR cohort). The converter domain regions identified in both the pre-stroke and post-stroke states showed enrichment of disease-associated variants in the replication dataset (pre-stroke p = 0.0019, post-stroke p = 1.7 x 10^−4^).

We extended our analysis of three-dimensional protein space to examine surface regions of β-cardiac myosin. To find such domains, we first defined surface-exposed amino acids by their accessibility to a spherical probe with a radius of 2.5 Å (the approximate size of an amino acid side chain) and approximated the surface distance between any two residues^23^ (see Methods and Supplementary Figure S3 and Fig. 4). The surface of β-cardiac myosin contains 568 of the 765 residues in the S1 domain (74%). Of these, 71 are associated with HCM (72% of all HCM variants) and 79 are found in a reference population (71% of all reference variants), suggesting that variants in both cohorts are evenly distributed between the surface and core of the protein (chi-square p=0.51). We then applied our spatial scan statistic to the surface of β-cardiac myosin. Using the myosin pre-stroke model, we identified a region of the surface covering 277 of the 568 surface amino acid residues (p = 0.002, Fig. 3a,b), including the converter domain and the mesa, that is highly enriched for disease-associated variation. Strikingly, the region contains 52 of the 71 surface HCM-associated missense variants and only 27 of the 79 surface non-disease associated missense variants (Fig. 3d), whereas the remainder of the myosin surface (Fig. 3c) covers 291 residues and contains only 19 disease-associated variants compared with 52 non-disease associated variants (Fig. 3d). The identified converter/mesa region was also enriched in the replication data set (p = 2.5 x 10^−5^).

**Fig 3.**
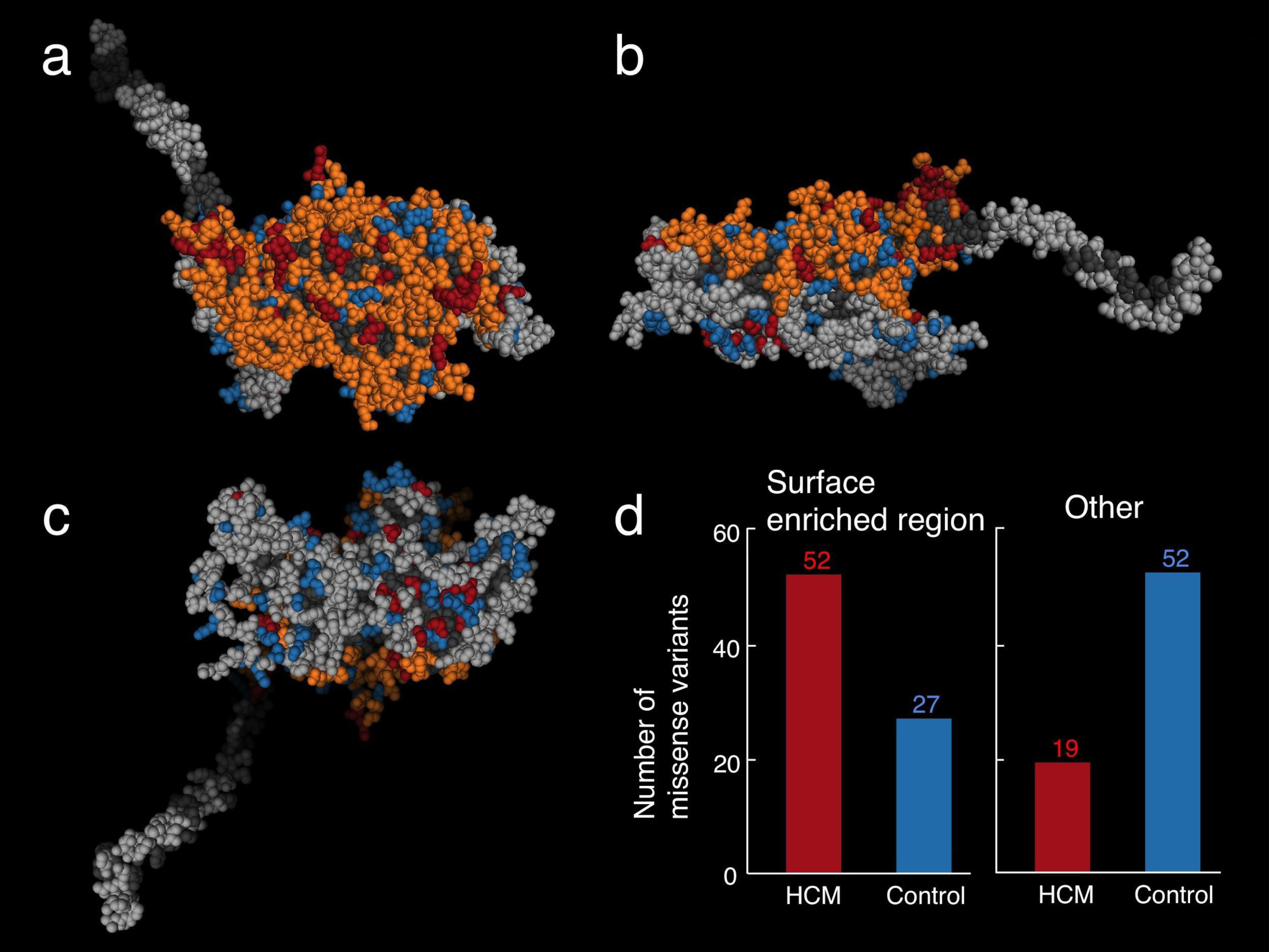
Surface spatial scan analysis identifies a larger surface region enriched for HCM-associated missense variation. (a) A similar view of the pre-stroke model to that in Figure 2b, looking directly down onto the mesa. The residues are colored as follows: orange - surface region enriched for HCM variants, blue - missense variants seen only in Exac, red - missense variants seen in SHaRe, light grey - residues considered to be on the surface, dark grey - residues not considered to be on the surface. The HCM-enriched surface region identified covers the entire mesa plus the adjoining converter domain. (b) The same side view of the pre-stroke S1 motor domain as shown Figures 1 and 2a. (c). A view of the pre-stroke model viewing the side opposite the mesa. This surface is not enriched for HCM variants. (d). The number of HCM-variants (SHaRe) and non-disease associated (ExAC) variants identified in the surface enriched region (left) and in the sum of all other parts of the myosin surface (right).

**Fig 4.**
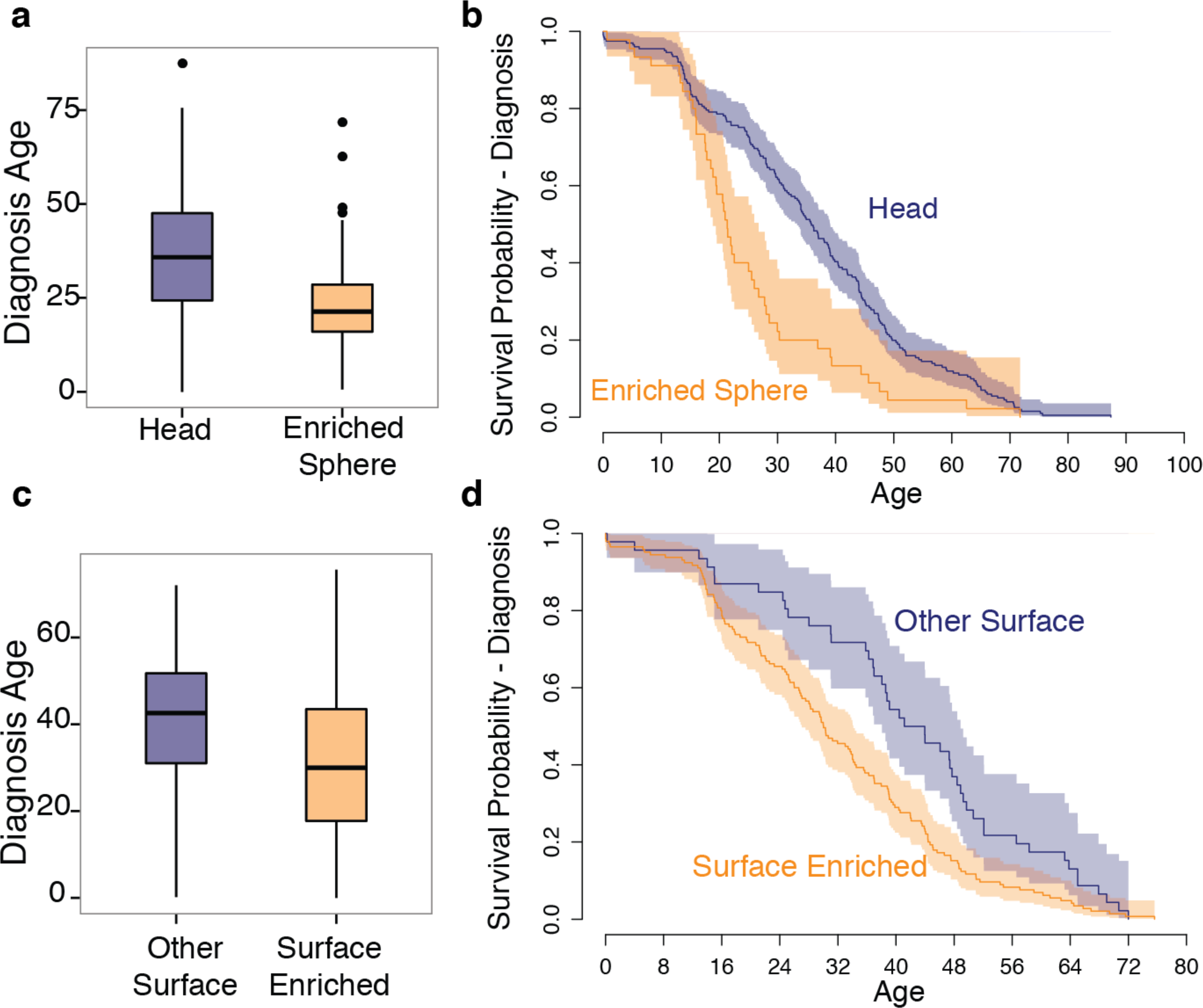
Comparison of clinical phenotypes between enriched regions and other regions in β-cardiac myosin. (a) Age of diagnosis of patients with HCM variants in the enriched spherical converter region (orange) compared with those in other parts of the myosin head (blue). (b) Kaplan-Meier curves of age at diagnosis compared between HCM variants in the enriched spherical converter region (orange) compared with those in other parts of the myosin head (blue). Shading indicates 95% confidence intervals for the survival curve. (c) Age of diagnosis of patients with HCM variants in the enriched surface region (orange) compared with those in other parts of the myosin head surface (blue). (d) Kaplan-Meier curves of age at diagnosis compared between HCM variants in the enriched surface region (orange) compared with those in other parts of the myosin head surface (blue). Shading indicates 95% confidence intervals for the survival curve.

Using the same procedure to search the surface regions of the myosin post-stroke structure, we detected a smaller enriched region of 122 amino acid residues again covering the converter but with a reduced portion of the myosin mesa (Supplementary Figure S5). During the myosin power stroke, the converter moves away from the mesa (compare Fig. 1a and Fig. 1b, Fig.3 and Supplementary Figure S5), so the enriched converter/mesa region is no longer contiguous and available for intra- or intermolecular interactions in the post-stroke state (Fig. 1b, Fig. 3a,b). Indeed, the enriched amino acid residues in the post-stroke model move significantly more in three-dimensional space between the pre and the post-stroke models (Wilcoxon p < 2 x 10^−16^) than other amino acid residues on the surface of the protein. These data suggest that myosin conformational changes during the actin-activated chemomechanical cycle may be important not only for transducing force, but also for modulating the size and shape of this surface region and altering its availability for binding. Indeed, the functional importance of the converter/mesa region is further supported by the presence of the binding site for omecamtiv mecarbil, a recently described small molecule modulator of cardiac myosin currently in clinical trials for the treatment of heart failure^24,25^.

Next, we tested the myosin S2 fragment for regions enriched for disease associated variation. The spatial scan analysis revealed that the first half of the S2 fragment is enriched for disease variants (p=0.003, Supplementary Figure S6). This proximal part of S2 has been shown to bind to the amino-terminal part of myosin binding protein C (MyBP-C)^26^, a sarcomere protein that is most frequently mutated in patients with HCM^14^. The enrichment of disease-associated variants in this region suggests that binding between myosin S2, MyBP-C (and potentially other partners) is important for development of HCM.

To further investigate the contribution of the genetically-constrained regions to disease, we compared the clinical features of patients with variants in these regions to patients with variants elsewhere in *MYH7*. The clinical profile of HCM is highly variable, with some patients living a normal lifespan with minimal symptoms and others dying suddenly or requiring cardiac transplantation at a young age^27^. Similarly, age at presentation of HCM varies widely between patients and earlier onset is correlated with a more severe phenotype^28^. We find that HCM patients with a variant inside the spherical enriched region are 11.2 years younger at diagnosis (24.9 vs. 36.1 years old, Wilcoxon p = 6.7 x 10^−5^) (Fig. 4a-b) than patients harboring other variants in the myosin head. The presence of a variant in the HCM-enriched surface region is associated with a 10.0-year earlier age at diagnosis (age 31.5 vs. age 41.5, Wilcoxon p = 1.6 x 10^−4^) than those with other surface variants (Fig. 4c-d). In addition, we find an increased hazard for clinical outcomes in the surface enriched region (HR = 1.918, p=0.023), though not in the spherical enriched region (Supplementary Figure S7-8, Supplementary Text 2). These findings demonstrate that analysis of genetic constraint in protein space can reveal domains with both increased disease burden and pathogenicity.

Our study demonstrates the power of integrating detailed structural information with large clinical and genetic databases to identify regions associated with functional importance and disease severity in Mendelian diseases. We find that variants associated with HCM are enriched in the β-cardiac myosin converter domain, where they lead to more severe outcomes. We provide the first evidence that similar clustering and pathogenicity are present in a surface spanning the converter domain and the recently-described mesa. Because amino acid residues forming the mesa come from disparate locations in the nucleotide sequence, discovery of this region depends on integration of protein structural information. The pronounced shift of the converter/mesa surface during the power stroke raises the mechanistic hypothesis that these variants exert their deleterious effect selectively in the pre-stroke state, perhaps by disrupting dynamic binding interactions. In summary, these findings highlight the importance of considering data from human genetics in the context of the dynamic, 3-dimensional protein structure, and illustrate a new approach to structure-function analysis in genetic diseases.

## Methods

### SHaRe Database

The Sarcomeric Human Cardiomyopathy Registry (SHaRe), a multicenter database that pools de-identified patient-level data from established institutional datasets at participating sites. At the time of analysis, the registry contained clinical and genetic testing data on 2,913 patients with HCM. Over 1,000 of these patients have pathogenic variants in *MYH7* and *MYBPC3*. This database contains individuals from 9 inherited disease centers throughout the world, including Brigham and Women’s Hospital, Children’s Hospital Boston, Erasmus Medical Center, Careggi University in Florence, Stanford Center for Inherited Cardiovascular Disease, University College London, University of Michigan, the Laboratory of Genetics and Molecular Cardiology in Sao Paolo, and Akureyri Hospital Iceland. The database includes demographic data, medical history, echocardiogram data, genetic testing results, and many other data relating to cardiac health and clinical outcomes.

### ExAC

The Exome Aggregation Consortium^13^ released data from 60,706 exomes from multiple sequenced cohorts that are not enriched for rare diseases such as HCM. We downloaded ExAC data for the canonical *MYH7* transcript ENST00000355349 on August 27, 2015.

### Variant Filtering and Inclusion Criteria

Variants in *MYH7* from the SHaRe database were filtered for quality purposes. Only exonic variants were included in the analysis. We included all exonic missense variants seen in HCM patients in clinical genetic testing. For comparison, we downloaded data from the *MYH7* gene from the Exome Aggregation Consortium (ExAC) on August 27, 2015. We excluded variants with a population specific frequency of greater than 1/2000 in the ExAC data, as these common variants are unlikely to be causal for HCM based upon the population prevalence of the disease. For the spatial scan and clinical outcomes analysis, we analyzed only missense variants.

For validation purposes, we also performed a subset of analyses including variants expertly classified as “pathogenic” or “likely pathogenic” while excluding variants of unknown significance. This is to ensure that variants of unknown significance (VUS) are not driving the results of the analyses. In addition, we also performed a validation using variants found in individuals of European ancestry from ExAC and individuals with a reported race of white, to ensure that global population structure was not confounding our analysis (Supplementary Text 3).

We generated an independent validation data set combining previously published analyses of HCM variants from other medical centers with 42,930 exomes from the DiscovEHR sequencing project involving the Regeneron Genetics Center and Geisinger Health System. Once again, we included only missense variants and removed variants with an allele frequency greater than 1 in 2000 in the DiscovEHR exomes (Supplementary Text 1).

### Development of human β-cardiac myosin protein models

We developed human β-cardiac myosin S1 models based on human motor domain structural data to best represent the human form of the cardiac myosin. We retrieved the protein sequence of human β-cardiac myosin and the human cardiac light chains from UNIPROT database^29^: myosin heavy chain motor domain (MYH7) - P12883, myosin essential light chain (MLC1) - P08590, and myosin regulatory light chain (MLC2) - P10916. We used a multi-template homology modeling approach to build the structural coordinates of MYH7 (residues 1-840), MCL1 (residues 1-195), MCL2 (residues 1-166) and S2 (residues 841-1280). We obtained the three dimensional structural model of S1 in the pre- and post- stroke states by integrating the known structural data from solved crystal structures, as described below.

Homology modeling of the pre-stroke structure was performed with template structures of the smooth muscle myosin motor domain^30^ (PDB id: 1BR1) and the scallop smooth muscle myosin light chain domain^31^ (PDB id: 3TS5). The templates used for the modeling of the post-stroke structure were obtained from the human β-cardiac myosin motor domain^32^ (PDB id: 4P7H) and the rigor structure from the squid myosin motor domain^33^ (PDB id: 3I5G). Missing regions in the myosin motor domain (loop1, loop2) were each separately built using the ModLoop program^34^and regions in the regulatory light chains that were not solved in the crystal structures were independently modelled using the I-TASSER prediction^35^ method, and they were used as individual templates. Sequence alignment between MHY7, MLC1, and MLC2 with their respective structural templates were obtained. The models of pre- and post-stroke structures were acquired using the MODELLER package^36^. We used a multi-template modeling method: 100 models were obtained and the best model was selected based on the DOPE score. The structural models were energy minimized using SYBYL7.2 (Tripos Inc.) to remove potential short contacts. The final three-dimensional models of the pre- and post-stroke structures were validated using RAMPAGE^37^, which provides a detailed check on the stereochemistry of the protein structure using the Ramachandran map.

The S2 region is a long coiled-coil structure; hence we used the template from the Myosinome database^38^. Modeling was done using the MODELLER package, selection of best model and validation of the model was done as described above. Visualizations were performed using PyMOL version 1.7.4 (http://www.pymol.org).

### Statistical Methods

Comparisons between the ExAC and SHaRe variant locations in *MYH7* were performed using the Kolmogorov-Smirnov test statistic. All statistical analyses were performed in R version 3.1.2 ^39^ and many graphs were prepared using ggplot2^40^.

### Spatial Scan Statistic

For the spatial scan analysis, we compared the locations of unique variants observed in HCM patients in SHaRe with the locations of variants observed in ExAC. The Spatial Scan Statistic exhaustively searches three-dimensional windows of a predefined set of sizes and shapes throughout the human β-cardiac myosin molecule for regions with an increased proportion of HCM-associated (SHaRe) variants. Let *p_w_* be the proportion of variation within a window that is HCM-associated, let *q_w_* be the proportion of variation outside the window which is HCM-associated. For each window, we calculate the binomial likelihood ratio statistic comparing the null model where *p_w_* and *q_w_* both equal the overall rate *r* against the model where *p_w_* is not equal to *q_w_*. The likelihood ratio statistic for each window is as follows:

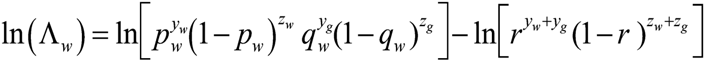

where *y_w_* is the number of HCM-associated variants within the window, *z_w_* is the number of other variants in the window, *y_g_* is the number of HCM-associated variants outside the window, *z_g_* is the number of other variants outside the window, and *r* is the overall proportion of HCM-associated variants. The test statistic is the maximum of the observed likelihood ratio statistics for all windows. Significance is assessed through permutation analysis by permuting the variant labels.

For the first analysis, we used spherical windows based upon the three-dimensional locations of amino acids in the human β-cardiac myosin molecule. The myosin S1 models include residues 1 to 841. We excluded the disordered loop regions between residues 205 and 211 and residues 627 and 640, as the positions of these amino acid residues are not well defined. We compared the observed missense variants in human β-cardiac myosin in SHaRe and in ExAC. We generated lists of unique missense variants from the SHaRe and ExAC datasets. A variant was considered HCM-associated if it was observed in a SHaRe patient diagnosed with HCM and not HCM associated if the variant was only observed in the ExAC data. Variants were assigned to three-dimensional locations in the human β-cardiac myosin model based upon the position of the alpha carbon atom of their corresponding amino acid residue. For each amino acid residue in the myosin S1, we tested spherical windows with radii of 10, 12.5, 15, 17.5, 20, 22.5, and 25 Angstroms centered on the alpha carbon for enrichment of HCM associated variants. The maximum test statistic for the entire set of windows in the model was calculated and significance was assessed through permutation of variant labels using 1000 permutations. For validation, we performed the analysis above removing all missense variants classified as Variants of Unknown Significance (Supplementary Text 3). We also performed the analysis above using only missense variants observed in the European population in ExAC or in European ancestry HCM cases.

### Surface Analysis

In addition to the spherical windows defined above, we define windows based upon the exposed surface of the human β-cardiac myosin molecule. For this analysis, we estimate the surface distance between any two amino acids. We developed the following procedure to perform this estimation. First, we calculated the solvent excluded surface for the human β-cardiac myosin models using the MSMS program^23^ with a 2.5 Å radius sphere as a probe, which approximates the size of amino acid side chain interactions. For the calculated surface, the MSMS program returns a large net of vertices of the surface, each connected to multiple other points on the surface. We use these vertices and connections to build a weighted graph of the surface of the molecule, with the vertices as nodes and the edges as connections weighted by the Euclidean distance between the two connected vertices. Then, for each amino acid, we assign it to the point on the surface that is the closest point, on average, to all of the non-hydrogen atoms of the amino acid. This average distance is used as an estimate of the depth of the amino acid. Amino acids with average depths of greater than 4 Angstroms for the surface model were considered to be not on the surface. We used the A-star algorithm to calculate the distance on the surface graph between two amino acid residues. In this analysis we included only amino acids in the ‘head’ of the human β-cardiac myosin molecule, defined as amino acids 1 to 784. The lever arm was excluded. We excluded the disordered loop regions between residues 205 and 211 and residues 627 and 640.

Based upon this set of pairwise calculated distances, we define planar surface regions of the MYH7 molecule for analysis as all the amino acids within a certain distance of any given ‘center’ amino acid. We exclude non-surface amino acids, based upon their estimated depth. We once again perform the spatial scan statistic as described above to identify surface regions of increased genetic burden.

### S2 Fragment Analysis

We performed the spherical spatial scan statistic as described above to test for enrichment in the S2 fragment of myosin. We used spherical window sizes of 10, 20, 30, 40, 50, 60, 70, 80, 90, and 100 Angstroms centered on the alpha carbon of each amino acid residue in the S2 model. For this analysis, we compared the region between amino acid residue 838 and amino acid residue 1112. The analysis was truncated at amino acid residue 1112 due to low sequencing coverage following residue 1112 in the ExAC data.

### Clinical phenotype analysis

We perform outcomes analysis using statistical methods implemented in R version 3.1.2. For the analysis of age at diagnosis only known probands were included. We compared the primary diagnosis ages using a Wilcoxon test. For the age at event analyses (Supplementary Text 2), the composite outcomes were defined as follows: The arrhythmic outcome consisted of cardiac arrest, ICD firing, and sudden cardiac death. The heart failure outcome combined the events of end-stage HCM (defined as the left ventricular ejection fraction falling below 55%), New York Heart Association Class III or IV status, transplant operation, and left ventricular assistive device implantation. The overall composite outcome combined the arrhythmic and heart failure outcomes as well as including the outcomes of atrial fibrillation, stroke, and death (all causes). Individuals were considered to enter the study at their diagnosis age and were censored at their last known age. We compared hazard ratios for each region using the Cox proportional hazards model adjusting for gender.

## Acknowledgements

The authors would like to thank Jonathan Fox for guidance during the early stages of the SHaRe registry and Aleks Pavlovic for support in obtaining and curating the clinical data.

## Competing interests

J.A.S is a founder of and owns shares in Cytokinetics, Inc. and MyoKardia, Inc., biotech companies that are developing therapeutics that target the sarcomere. E.M.G is an employee and owns shares in MyoKardia, Inc. E.A.A is a founder of Personalis, Inc. C.D.B. is on the Scientific Advisory Boards of http://Ancestry.com, Personalis, Liberty Biosecurity, and Etalon DX. C.D.B. is also a founder and chair of the SAB of IdentifyGenomics.

## References

1 Krendel, M. & Mooseker, M.S. Myosins: tails (and heads) of functional diversity. Physiology 20, 239–251 (2005).

2 Ruppel, K.M. & Spudich, J.A. Structure-function analysis of the motor domain of myosin. Annu. Rev. Cell Dev. Biol. 12, 543–573 (1996).

3 Rall, J.A. Mechanism of Muscular Contraction.(2014).

4 Oldfors, A. Hereditary myosin myopathies. Neuromuscul. Disord. 17, 355–367 (2007).

5 Maron, B.J., Casey, S.A., Hauser, R.G. & Aeppli, D.M. Clinical course of hypertrophiccardiomyopathy with survival to advanced age. J. Am. Coll. Cardiol. 42, 882–888 (2003).

6 Walsh, R., Rutland, C., Thomas, R. & Loughna, S. Cardiomyopathy: a systematic review of disease-causing mutations in myosin heavy chain 7 and their phenotypic manifestations. Cardiology 115, 49–60 (2010).

7 Jellis, C.L. & Desai, M.Y. Hypertrophic cardiomyopathy: still connecting the dots between genotype and phenotype. Cardiovasc Diagn Ther 5, 156–159 (2015).

8 Pan, S. et al Cardiac Structural and Sarcomere Genes Associated with Cardiomyopathy Exhibit Marked Intolerance of Genetic Variation. Circ. Cardiovasc. Genet. 602–610 (2012).

9 Geisterfer-Lowrance, A. a. et al A molecular basis for familial hypertrophic cardiomyopathy: a beta cardiac myosin heavy chain gene missense mutation. Cell 62, 999–1006 (1990).

10 Buvoli, M., Hamady, M., Leinwand, L.A. & Knight, R. Bioinformatics assessment of beta-myosin mutations reveals myosin’s high sensitivity to mutations. Trends Cardiovasc. Med. 18, 141–149 (2008).

11 Kapplinger, J.D. et al Distinguishing hypertrophic cardiomyopathy-associated mutations from background genetic noise. J. Cardiovasc. Transl. Res. 7, 347–361 (2014).

12 Examining Prevailing Genotype-Phenotype Correlations in Hypertrophic Cardiomyopathy: Findings From The Sarcomeric Human Cardiomyopathy Registry (SHaRe). Circulation 132, 2267–2285 (2015).

13 Lek, M. et al Analysis of protein-coding genetic variation in 60,706 humans. bioRxiv (2015). doi:10.1101/030338

14 Alfares, A.A. et al Original Research Article Results of clinical genetic testing of 2,912 probands with hypertrophic cardiomyopathy: expanded panels offer limited additional sensitivity. (2015). doi:10.1038/gim.2014.205

15 Spudich, J.A. The myosin mesa and a possible unifying hypothesis for the molecular basis of human hypertrophic cardiomyopathy. Biochemical Society Transactions 43, 64–72 (2015).

16 Kulldorff, M. & Martin, K. A spatial scan statistic. Communications in Statistics - Theory and Methods 26, 1481–1496 (1997).

17 Ionita-Laza, I., Makarov, V., ARRA Autism Sequencing Consortium & Buxbaum, J.D. Scan-statistic approach identifies clusters of rare disease variants in LRP2, a gene linked and associated with autism spectrum disorders, in three datasets. Am. J. Hum. Genet. 90, 1002–1013 (2012).

18 Seebohm, B. et al Cardiomyopathy mutations reveal variable region of myosin converter as major element of cross-bridge compliance. Biophys. J. 97, 806–824 (2009).

19 Köhler, J. et al Mutation of the myosin converter domain alters cross-bridge elasticity. Proc. Natl. Acad. Sci. U. S. A. 99, 3557–3562 (2002).

20 Swank, D.M. et al The myosin converter domain modulates muscle performance. Nat. Cell Biol. 4, 312–316 (2002).

21 Garcia-Giustiniani, D. et al Phenotype and prognostic correlations of the converter region mutations affecting the myosin heavy chain. Heart 101, 1047–1053 (2015).

22 Woo, A. Mutations of the myosin heavy chain gene in hypertrophic cardiomyopathy: critical functional sites determine prognosis. Heart 89, 1179–1185 (2003).

23 Sanner, M.F., Olson, A.J. & Spehner, J.C. Reduced surface: an efficient way to compute molecular surfaces. Biopolymers 38, 305–320 (1996).

24 Malik, F.I. et al Cardiac myosin activation: a potential therapeutic approach for systolic heart failure. Science 331, 1439–1443 (2011).

25 Winkelmann, D.A., Forgacs, E., Miller, M.T. & Stock, A.M. Structural basis for drug-induced allosteric changes to human β-cardiac myosin motor activity. Nat. Commun. 6, 7974 (2015).

26 Starr, R. & Offer, G. The interaction of C-protein with heavy meromyosin and subfragment-2. Biochem. J 171, 813–816 (1978).

27 Maron, B.J. et al Clinical Course of Hypertrophic Cardiomyopathy in a Regional United States Cohort. JAMA 281, 650 (1999).

28 Maron, B.J. et al Hypertrophic Cardiomyopathy in Adulthood Associated With Low Cardiovascular Mortality With Contemporary Management Strategies. J. Am. Coll. Cardiol. 65, 1915–1928 (2015).

29 UniProt Consortium. Update on activities at the Universal Protein Resource (UniProt) in 2013. Nucleic Acids Res. 41, D43–7 (2013).

30 Dominguez, R., Freyzon, Y., Trybus, K.M. & Cohen, C. Crystal structure of a vertebrate smooth muscle myosin motor domain and its complex with the essential light chain: visualization of the prepower stroke state. Cell 94, 559–571 (1998).

31 Kumar, V. S. S. et al Crystal structure of a phosphorylated light chain domain of scallop smooth-muscle myosin. Biophys. J. 101, 2185–2189 (2011).

32 Winkelmann, D.A., Miller, M.T., Stock, A.M. & Liu, L. Structure of Human beta-Cardiac Myosin Motor Domain at 3.2 A. Mol. Biol. Cell 4705 (2011).

33 Yang, Y. et al Rigor-like structures from muscle myosins reveal key mechanical elements in the transduction pathways of this allosteric motor. Structure 15, 553–564 (2007).

34 Fiser, A. & Sali, A. ModLoop: automated modeling of loops in protein structures. Bioinformatics 19, 2500–2501 (2003).

35 Yang, J. et al The I-TASSER Suite: protein structure and function prediction. Nat. Methods 12, 7–8 (2015).

36 Sali, A. & Blundell, T.L. Comparative protein modelling by satisfaction of spatial restraints. J. Mol. Biol. 234, 779–815 (1993).

37 Lovell, S.C. et al Structure validation by Cα geometry: ϕ,ψ and Cβ deviation. Proteins: Struct. Funct. Bioinf. 50, 437–450 (2003).

38 Syamaladevi, D.P. et al Myosinome: a database of myosins from select eukaryotic genomes to facilitate analysis of sequence-structure-function relationships. Bioinform. Biol. Insights 6, 247–254 (2012).

39 R Core Team. R: A Language and Environment for Statistical Computing. (2014). at http://www.R-project.org/

40 Wickham, H. ggplot2: Elegant Graphics for Data Analysis. (Springer Science & Business Media, 2009).

41 Blair, E., Price, S.J., Baty, C.J., Ostman-Smith, I. & Watkins, H. Mutations in cis can confound genotype-phenotype correlations in hypertrophic cardiomyopathy. J. Med. Genet. 38, 385–388 (2001).

42 Bos, J.M. et al Characterization of a phenotype-based genetic test prediction score for unrelated patients with hypertrophic cardiomyopathy. Mayo Clin. Proc. 89, 727–737 (2014).

43 Ingles, J. et al Compound and double mutations in patients with hypertrophic cardiomyopathy: implications for genetic testing and counselling. J. Med. Genet. 42, e59 (2005).

44 Kubo, T. et al Prevalence, clinical significance, and genetic basis of hypertrophic cardiomyopathy with restrictive phenotype. J. Am. Coll. Cardiol. 49, 2419–2426 (2007).

45 Millat, G., Gilles, M., Valérie, C., Hervé, C. & Robert, R. Development of a high resolution melting method for the detection of genetic variations in hypertrophic cardiomyopathy. Clin. Chim. Acta 411, 1983–1991 (2010).

46 Otsuka, H. et al Prevalence and distribution of sarcomeric gene mutations in Japanese patients with familial hypertrophic cardiomyopathy. Circ. J. 76, 453–461 (2012).

47 Richard, P. et al Hypertrophic cardiomyopathy: Distribution of disease genes, spectrum of mutations, and implications for a molecular diagnosis strategy. Circulation 107, 2227–2232 (2003).

48 Roncarati, R. et al Unexpectedly low mutation rates in beta-myosin heavy chain and cardiac myosin binding protein genes in Italian patients with hypertrophic cardiomyopathy. J. Cell. Physiol. 226, 2894–2900 (2011).

49 Song, L. et al Mutations profile in Chinese patients with hypertrophic cardiomyopathy. Clin. Chim. Acta 351, 209–216 (2005).

50 Wang, J. et al Malignant effects of multiple rare variants in sarcomere genes on the prognosis of patients with hypertrophic cardiomyopathy. Eur. J. Heart Fail. 16, 950–957 (2014).

51 Yu, B. et al Denaturing high performance liquid chromatography: high throughput mutation screening in familial hypertrophic cardiomyopathy and SNP genotyping in motor neurone disease. J. Clin. Pathol. 58, 479–485 (2005).

